# Source-to-Tap Investigation of the Occurrence of Nontuberculous Mycobacteria in a Full-Scale Chloraminated Drinking Water System

**DOI:** 10.1101/2024.04.18.590135

**Authors:** Katherine S. Dowdell, Sarah C. Potgieter, Kirk Olsen, Soojung Lee, Matthew Vedrin, Lindsay J. Caverly, John J. LiPuma, Lutgarde Raskin

## Abstract

Nontuberculous mycobacteria (NTM) in drinking water are a significant public health concern. However, an incomplete understanding of the factors that influence the occurrence of NTM in drinking water limits our ability to characterize risk and prevent infection. This study sought to evaluate the influence of season and water treatment, distribution, and stagnation on NTM in drinking water. Samples were collected source-to-tap in a full-scale, chloraminated drinking water system approximately monthly from December 2019 to November 2020. NTM were characterized using culture-dependent (plate culture with matrix-assisted laser desorption ionization-time of flight mass spectrometry [MALDI-TOF MS] isolate analysis) and culture-independent methods (quantitative PCR and genome resolved metagenomics). Sampling locations included source waters, three locations within the treatment plant, and five buildings receiving water from the distribution system. Building plumbing samples consisted of first draw, five-minute flush, and full flush cold-water samples. As the study took place during the COVID-19 pandemic, the influence of reduced water usage in three of the five buildings was also investigated. The highest concentrations of NTM source-to-tap were found in the summer first draw building water samples (10^7^ gene copies/L), which also had the lowest chloramine concentrations. Flushing was found to be effective for reducing NTM and restoring disinfectant residuals, though flush times necessary to improve water quality varied by building. Clinically-relevant NTM species, including *Mycobacterium avium*, were recovered via plate culture, with increased occurrence observed in buildings with higher water age. Four of five NTM metagenome-assembled genomes were identified to the species level and matched identified isolates.

**Importance:** NTM infections are increasing in prevalence, difficult to treat, and associated with high mortality rates. Our lack of understanding of the factors that influence NTM occurrence in drinking water limits our ability to prevent infections, accurately characterize risk, and focus remediation efforts. In this study, we comprehensively evaluated NTM in a full-scale drinking water system, showing that various steps in treatment and distribution influence NTM presence. Stagnant building water contained the highest NTM concentrations source-to-tap and was associated with low disinfectant residuals. We illustrated the differences in NTM detection and characterization obtained from culture-based and culture-independent methods, highlighting the complementarity between these approaches. We demonstrated that focusing NTM mitigation efforts in building plumbing systems, which have the highest NTM concentrations source-to-tap, has potential for immediate positive effects. We also identified steps during treatment that increase NTM levels, which provides beneficial information for utilities seeking to reduce NTM in finished water.

## Introduction

Nontuberculous mycobacteria (NTM) respiratory infections are a significant and growing health concern, with numerous recent studies reporting increasing prevalence (1–3). The impact of NTM infections is illustrated in high associated mortality rates and healthcare costs compared to infections by enteric waterborne pathogens (4–6). Though the sources of NTM infections are often difficult to determine, drinking water exposure is believed to be a major route of infection (7, 8). However, the factors that shape NTM concentrations at the tap are poorly understood. This limitation is due to several reasons, including the complexity of quantification methods, the lack of regulations governing NTM in drinking water, and the difficulty in linking specific environmental sources to infections (9). A greater understanding of NTM occurrence, species composition of NTM populations, and factors that contribute to higher NTM levels in drinking water is crucial for the development of quantitative risk assessments and risk mitigation strategies (10, 11).

Building plumbing characteristics have been shown to greatly influence tap water quality. Building plumbing properties, including high surface-to-volume ratios, periods of stagnation, and equilibration with building temperatures, create an environment that contributes to the growth of biofilms and high concentrations of certain opportunistic human pathogens (OPs), including NTM (12, 13). Although previous studies have found that building plumbing may contribute to high concentrations of NTM, few studies have evaluated the impacts of flushing, seasonal variations, and water quality parameters (14–17). Previous studies have found concentrations of NTM in building plumbing as high as 10^5^ - 10^7^ gene copies per liter (15, 18–20). The distribution system has also been shown to influence NTM, with higher concentrations of NTM or greater occurrence of clinically-relevant species linked to higher distribution system residence times (14, 21). Source water type (22) and drinking water treatment processes may also influence the NTM entering the distribution system, though few studies have investigated these topics in full-scale systems (23–27). Another factor linked to NTM occurrence is the type of secondary disinfectant used, with several studies reporting increased rates of NTM detection or higher NTM concentrations in distribution systems using monochloramine (22, 28–30). The COVID-19 pandemic brought additional concerns about OPs in building water due to low water use (31). Although most studies focused only on *Legionella pneumophila* (32–34), one study reported *Mycobacterium avium* complex up to 10^5^ gene copies per liter in stagnant building plumbing water (16).

Culture-based methods have long been the primary means for detecting NTM in environmental samples. Culture-based methods are often favored because they detect viable bacteria that can be used for additional analyses. However, existing culture-based methods for recovery of NTM from drinking water lack standardization, are laborious, and, despite the use of decontamination or antibiotic-containing media, are often limited in their specificity for NTM, meaning that additional methods are required to confirm species identity (35–37). Further, species- and strain-level variation in susceptibility to decontamination treatments and antibiotics may introduce bias in quantifying NTM populations in mixed communities in environmental samples (36–38). Quantitative PCR (qPCR) is often favored by researchers for quantifying NTM because it is rapid and typically highly specific at the genus or species level (39–43). However, unless viability pre-treatments are used (e.g. propidium monoazide or ethidium monoazide) (44–46), qPCR does not provide information regarding viability. Recently, amplicon sequencing and shotgun metagenomics have been used to determine the relative abundance of NTM in microbial communities, with some studies reporting the enrichment of NTM in building plumbing biofilms (14, 47–49). However, DNA sequencing methods may fail to detect low abundance microorganisms like NTM, they do not show viability, and the PCR employed in amplicon sequencing introduces bias (50, 51). Additionally, sequencing methods can only yield relative, rather than absolute abundances unless quantitative metagenomics is employed (52, 53). NTM are also difficult to lyse, which can impact DNA recovery and NTM detection (54). As methods for NTM quantification are not standardized, cross-study comparisons are challenging, particularly between studies that use culture-based methods versus those that use culture-independent methods.

The risk to human health associated with NTM in drinking water is also dependent on the species present. Species within *Mycobacterium* range from non-pathogenic, to OPs (e.g., *M. avium* and *Mycobacterium abscessus*), to non-environmental pathogens (e.g., *Mycobacterium tuberculosis*). Although the risk of infection due to OPs is highly dependent on the susceptibility of the host, it is estimated that approximately one dozen of the nearly 200 species of NTM are capable of causing human infection (55). Additionally, regional differences have been observed regarding which NTM species are most likely to cause human infection. Such regional differences could result in underestimating risk if the most appropriate species are not targeted (56, 57). Due to the difficulties in identifying NTM to the species- or complex-level, drinking water studies often quantify total NTM, with many targeting the ATP synthase subunit c (*atpE*) gene (15, 58–60), or only investigating one or a few species of concern, such as *M. avium* or *Mycobacterium intracellulare* (19, 22, 61, 62). Identification of NTM to the species level in colonies from plate culture is typically done using PCR and Sanger sequencing targeting genes such as β-subunit of RNA polymerase (*rpoB)* or heat shock protein 65 (*hsp65*) (63–66). Another alternative for colony identification is matrix-assisted laser desorption ionization-time of flight mass spectrometry (MALDI-TOF MS), which is a method that analyzes proteins and generates spectra that are matched to a spectral database (67, 68). Advantages of MALDI-TOF MS include that it is rapid, accurate, and commonly used in clinical laboratories (69, 70). Culture-independent methods for species-level identification of NTM typically employ short-read or long-read amplicon sequencing, with targets such as the *rpoB* or *hsp65* gene (14, 47, 71).

Although previous studies have quantified NTM at various locations in drinking water systems (14, 15, 28, 30, 72), few have conducted full, source-to-tap assessments (73). This study sought to characterize the factors that shape NTM concentrations at various stages of treatment and distribution, including impacts of source water selection, individual treatment processes, distribution, and stagnation in building plumbing. To this end, samples were collected approximately monthly over one year in a full-scale chloraminated system. Samples were collected from source waters (river and well water), through the drinking water treatment plant (WTP), and in five buildings (Sites A-E) served by the WTP. The estimated distribution system water ages for the five buildings ranged from approximately 16 hours at Site A to approximately 68 hours at Site E. Building plumbing samples included first draw, five-minute flush, and full flush cold-water samples. Samples were analyzed for routine physicochemical parameters, heterotrophic plate counts (HPC), and the presence of NTM, which were characterized using plate culture with MALDI-TOF MS for colony identification, qPCR, and genome resolved metagenomics. The onset of the COVID-19 pandemic and associated building closures occurred during the study, impacting water use at three of the buildings (Sites A, B, and E), facilitating an investigation of the impact of extended low water use on NTM and building water quality. The influence of stagnation was investigated through the collection of samples from buildings with various levels of flushing.

## Results

### NTM plate culture and MALDI-TOF MS

Presumptive NTM plate culture results ranged from less than the limit of detection (LOD) of 1 colony forming unit (CFU) per volume to 7.3×10^4^ CFU/L (Figure S1). LOD values ranged from 1 CFU/L to 333 CFU/L based on the sample volume. Median results in the source waters and through the WTP ranged from 1 CFU/L in the ozone effluent (n=7) to 48 CFU/L in the filter effluent (n=8). In the finished water, the median was 4 CFU/L (n=12). The highest results occurred in the first draw samples at Sites A (median: 1.2×10^4^ CFU/L, n=7) and B (median: 1.2×10^4^ CFU/L, n=7). The median first draw results in the other buildings ranged from 51 CFU/L at Site D (n=6) to 62 CFU/L at Site C (n=7). Five-minute flush and full flush sample median results ranged from 8 CFU/L in the Site E five-minute flush samples (n=8) to 2.4×10^2^ CFU/L in the Site B five-minute flush (n=7).

A subset of the colonies from the NTM culture plates (n=322), representing 47 of the samples, were identified using MALDI-TOF MS. Of these isolates, 60% (n=194) were identified as *Mycobacterium*, 34% (n=108) were not identified, indicating that they were other bacterial or fungal species not represented in the available spectral databases, and 6% (n=20) were identified as other genera including *Bacillus*, *Paenibacillus*, *Brevibacillus*, and *Micromonospora*, which all consist of endospore-forming bacterial species (Table 1) (74–77). Within the isolates identified as NTM, ten species or groups were identified, including *Mycobacterium arupense*, *Mycobacterium asiaticum*, *Mycobacterium aurum*, *M. avium*, *Mycobacterium chelonae* complex, *Mycobacterium franklinii*, *Mycobacterium gordonae*, *Mycobacterium llatzerense*, *Mycobacterium mucogenicum/ phocaicum* group, and *Mycobacterium peregrinum*. Of the recovered NTM species, *M. avium* and *M. chelonae* complex are of particular clinical relevance based on prevalence of infections (55, 78, 79).

**Table 1.** Results of matrix-assisted laser desorption ionization – time of flight mass spectrometry (MALDI-TOF MS) analysis of the plate culture isolates.

| Category | Identification | Number of Isolates (% of Total) | Score Range (Median) |
| --- | --- | --- | --- |
| *Mycobacterium* | *Mycobacterium arupense* | 2 (<1%) | 1.92 – 1.96 |
|  | *Mycobacterium asiaticum* | 1 (<1%) | 1.61 |
|  | *Mycobacterium aurum* | 4 (1%) | 1.98 – 2.27 (2.13) |
|  | *Mycobacterium avium* | 8 (2%) | 1.99 – 2.20 (2.09) |
|  | *Mycobacterium chelonae* complex | 84 (26%) | 1.62 – 2.12 (1.88) |
|  | *Mycobacterium franklinii* | 19 (6%) | 1.60 – 1.98 (1.77) |
|  | *Mycobacterium gordonae* | 2 (<1%) | 1.81 – 2.01 |
|  | *Mycobacterium llatzerense* | 1 (<1%) | 1.84 |
|  | *Mycobacterium mucogenicum/phocaicum* group | 65 (20%) | 1.82 – 2.39 (2.11) |
|  | *Mycobacterium peregrinum* | 8 (2%) | 2.10 – 2.31 (2.27) |
| Non-NTM | *Bacillus* spp. | 11 (3%) | 1.70 – 2.34 (1.94) |
|  | *Brevibacillus* spp. | 6 (2%) | 1.77 – 2.40 (1.96) |
|  | *Micromonospora* sp. | 1 (<1%) | 1.73 |
|  | *Paenibacillus* spp. | 2 (<1%) | 1.82 – 2.00 |
| Not identified | -- | 108 (34%) | 0.93 – 1.68 (1.21) |

MALDI-TOF MS results showed that none of the isolates recovered from the source waters and WTP ozone effluent and selected for identification were *Mycobacterium* spp. (Figure 1). In the river samples, River-Source isolates included three *Bacillus* spp. isolates and one isolate that could not be identified. For the River-Plant isolates, a larger fraction of isolates could not be identified (55%, n=10), and the remaining isolates were identified as *Bacillus* spp. (28%, n=5), *Paenibacillus* spp. (11%, n=2), and *Micromonospora* sp. (6%, n=1). All isolates from the well samples (Well-Source and Well-Plant) were microorganisms that could not be identified (n=46). Of the two isolates from the ozone effluent samples, one could not be identified, and the other was identified as a *Bacillus* sp.

**Figure 1.**
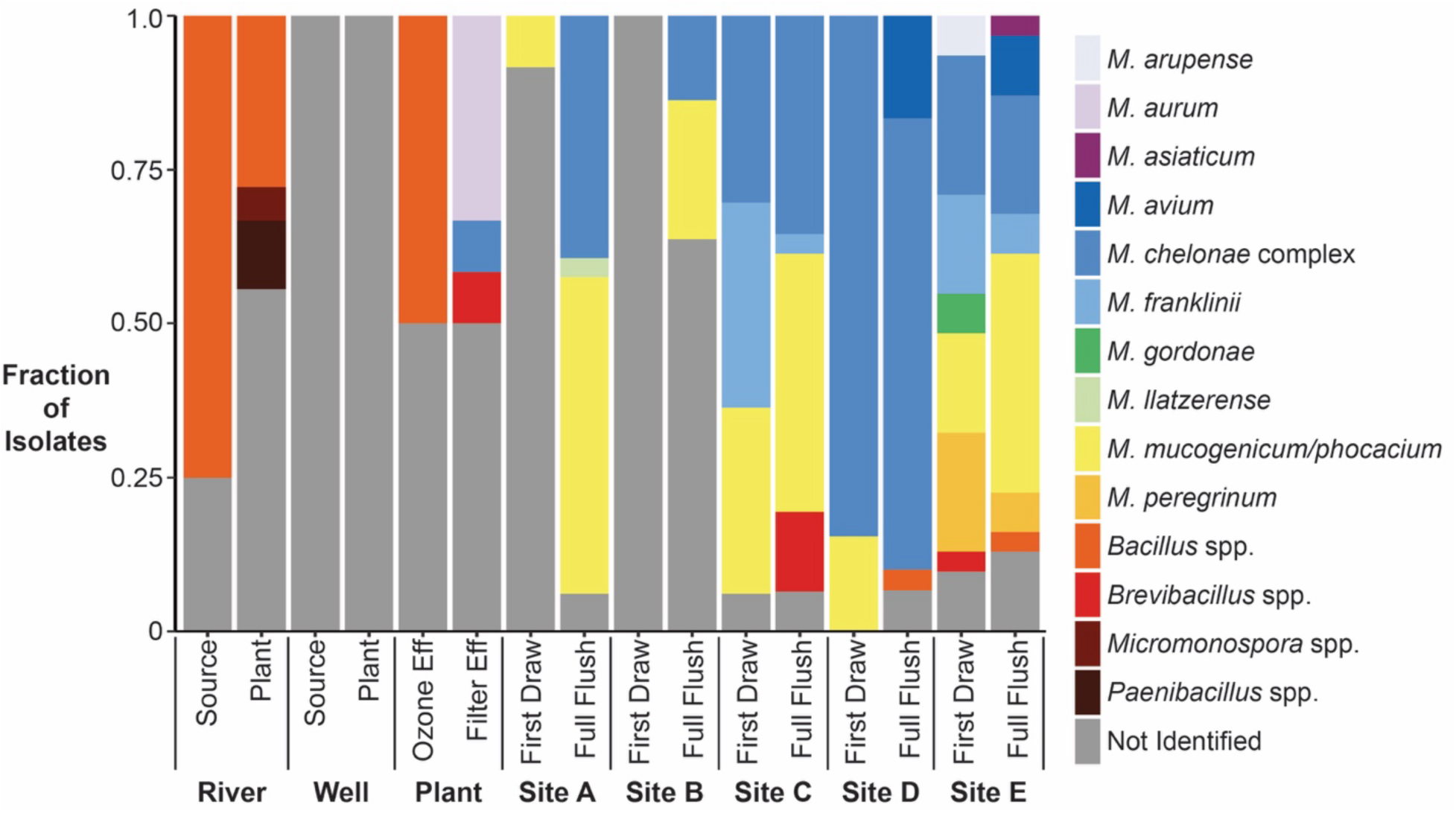
Results of matrix-assisted laser desorption ionization – time of flight mass spectrometry (MALDI-TOF MS) analysis for isolates from the NTM culture plates. No finished water isolates were recovered for the months analyzed. Not Identified: Sample spectra did not match spectral databases.

NTM isolates were identified in the WTP filter effluent and all downstream sampling locations, except the first draw samples from Site B. In the filter effluent, isolates included *M. aurum* (n=4), *M. chelonae* complex (n=1), *Brevibacillus* sp. (n=1), and six isolates that could not be identified. No isolates from the finished water were analyzed using MALDI-TOF MS due to the low number of isolates recovered. In the distribution system, the majority of isolates (68%, n=28 of 41) were NTM, with *M. chelonae* complex (n=8) and *M. mucogenicum/phocaicum* group (n=8) being the most frequently recovered. At Sites A and B, the fractions of NTM isolates recovered from first draw samples were low (<10%) but increased in the full flush samples (>35%). Increased recovery and diversity of NTM were observed at higher water age sites. At Sites D and E, the fractions of isolates identified as NTM were 93% (n=40 of 43) and 85% (n=53 of 62), respectively. The highest diversity in NTM isolates was observed at Site E, where eight different NTM species or groups were identified (*M. arupense*, *M. asiaticum, M. avium*, *M. chelonae* complex, *M. franklinii*, *M. gordonae*, *M. mucogenicum/phocaicum* group, and *M. peregrinum*). *M. avium* was only isolated from the full flush samples from Site D (n=5) and Site E (n=3).

The fraction of isolates identified as NTM for each sample analyzed using MALDI-TOF MS was used to adjust the presumptive NTM plate culture results, yielding the adjusted plate counts (Figure S2). Although the highest presumptive CFU/L results occurred in the first draw samples from Sites A and B, adjusted NTM CFU/L results were lower due to the large fraction of non-NTM isolates. The sample with the highest adjusted NTM plate count was the July Site C first draw sample at 5.6×10^2^ CFU/L. As most isolates picked from the Site C, D, and E culture plates were identified as NTM, the adjusted plate counts were generally similar to the presumptive NTM counts.

### NTM quantification using qPCR

Similar to the plate culture results, the highest concentrations of NTM using qPCR were observed in the first draw samples from Sites A and B (Figure 2). The lowest NTM gene copy concentrations were found in the Well-Source and the Ozone Effluent, where most samples were below the limit of quantification (LOQ) of 41 gene copies per reaction (gc/rxn). LOQs in gene copies per liter (gc/L) ranged from 2×10^2^ to 3×10^5^ gc/L and varied based on the sample volume and qPCR dilution factor. NTM gene copy concentrations in the river samples were higher, with a median of 7.3×10^4^ gc/L (n=11). Gene copy concentrations increased after filtration, from less than the LOQ in the Ozone Effluent to a median of 1.1×10^4^ gc/L in the Filter Effluent (n=7). In the Finished Water samples, the median NTM gene copy concentration was 1.4×10^3^ gc/L (n=8). At Sites A and B, maximum NTM gene copy concentrations in the first draw samples reached 5.5×10^5^ gc/L and 4.3×10^7^ gc/L, respectively.

**Figure 2.**
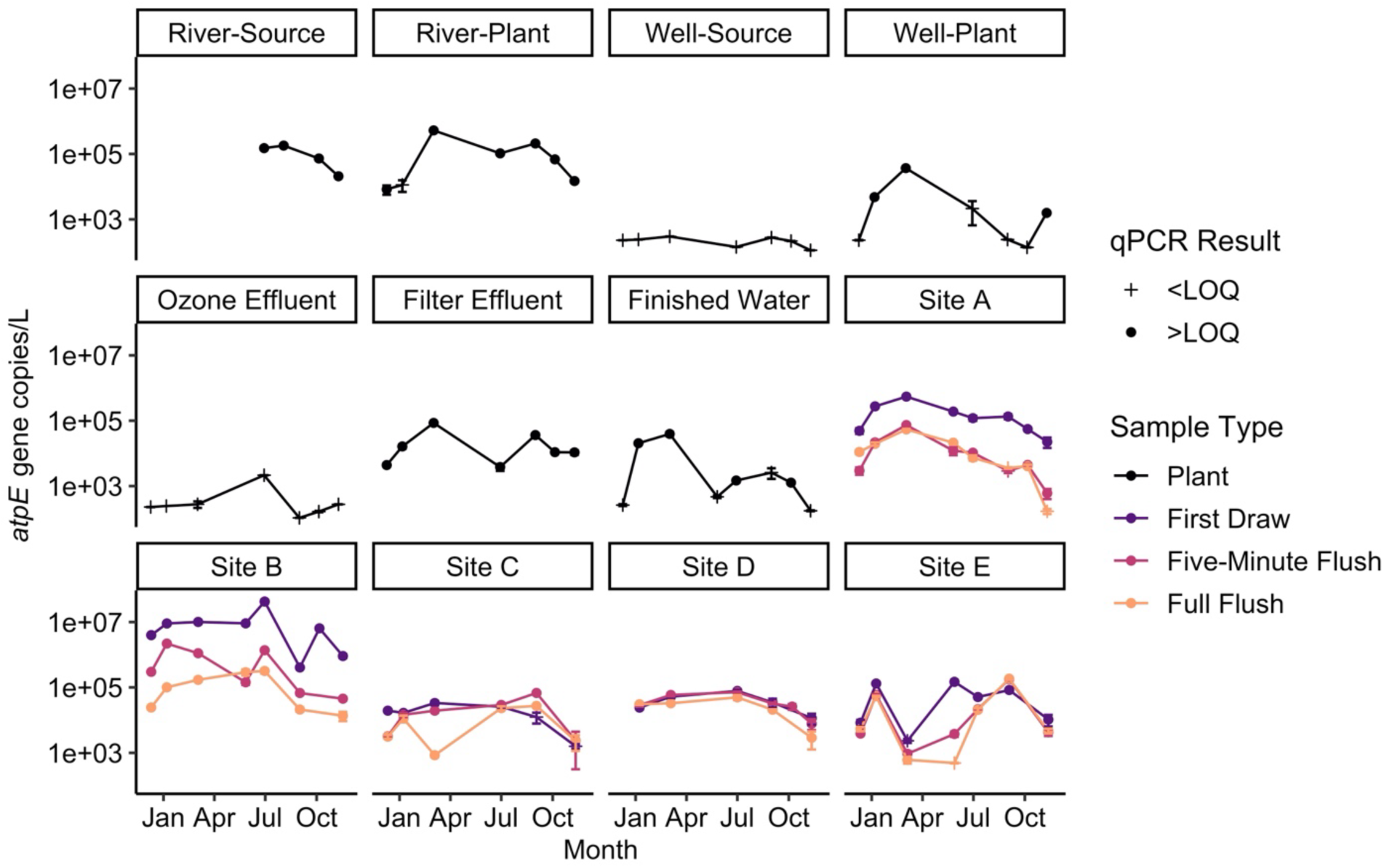
NTM *atpE* gene copy concentrations in all samples analyzed over the study period. Values below the LOQ (crosses) were set to one-half the LOQ and then converted to gene copies per liter. Values above the LOQ are shown as circles. Error bars show one standard deviation above and below the mean of triplicate qPCR reactions. Source: Based on data from Dowdell et al., 2022 (43).

In the building plumbing samples, the highest NTM gene copy concentrations were observed in the first draw samples (median: 6.7×10^4^ gc/L, n=34), followed by the five-minute flush samples (median: 2.4×10^4^ gc/L, n=34) and full flush samples (median: 2.1×10^4^ gc/L, n=33). NTM gene copy concentrations in the full flush samples, which captured distribution system water quality at the building locations, ranged from a median of 5.3×10^3^ gc/L at Site E to 1.0×10^5^ gc/L at Site B. Differences in NTM gene copy concentrations were observed over the sampling months. The months with the lowest median NTM gene copy concentrations in first draw samples were November (1.1×10^4^ gc/L, n=5) and December (3.4 ×10^4^ gc/L, n=4), while the highest median NTM gene copy concentrations in the first draw samples were found in October (3.3×10^6^ gc/L, n=2), May (1.9×10^5^ gc/L, n=3), and January (1.3×10^5^ gc/L, n=5). In the full flush samples, the months with the highest median NTM gene copy concentrations were March (3.3×10^4^ gc/L, n=5) and January (3.0×10^4^ gc/L, n=5), and the month with the lowest median NTM gene copy concentration was November (2.9×10^3^ gc/L, n=5).

### NTM species identified by metagenomic analysis

Processing of the metagenomic sequences from the WTP filter effluent, finished water, and full flush samples yielded five high quality (>50% completeness and <10% redundancy) NTM metagenome-assembled genomes (MAGs) (Figure 3, Table S4). Of these, four were identified to the species level, representing *M. phocaicum*, *M. llatzerense*, *M. gordonae,* and *M. arupense*. The MAG with the highest relative abundance across all samples was the one identified as *M. gordonae*, which occurred at a relative abundance of 0.91 reads per kilobase million (RPKM) in the December Site B full flush sample. The *M. gordonae* MAG was also detected in the Site B full flush samples from the other months, with relative abundances ranging from 0.08 to 0.51 RPKM, and in the October Site E full flush sample (0.15 RPKM). The NTM MAG was detected in the full flush samples from Site A in December, Site B in March, July, and October, and Site E in October, with relative abundances ranging from 0.04 to 0.14 RPKM. The only detection of the *M. phocaicum* MAG was in the December Site B full flush sample (0.49 RPKM). The highest relative abundance of the *M. arupense* MAG was in the July Site B full flush sample (0.65 RPKM). The *M. arupense* MAG was detected in the other Site B samples, as well as the Site A samples from December and October and the Site E samples from July and October. The *M. llatzerense* MAG was detected in the December Site B full flush sample (0.36 RPKM) and October Site E full flush sample (0.17 RPKM). No reads corresponding to the filter effluent or finished water samples met the 25% coverage minimum.

**Figure 3.**
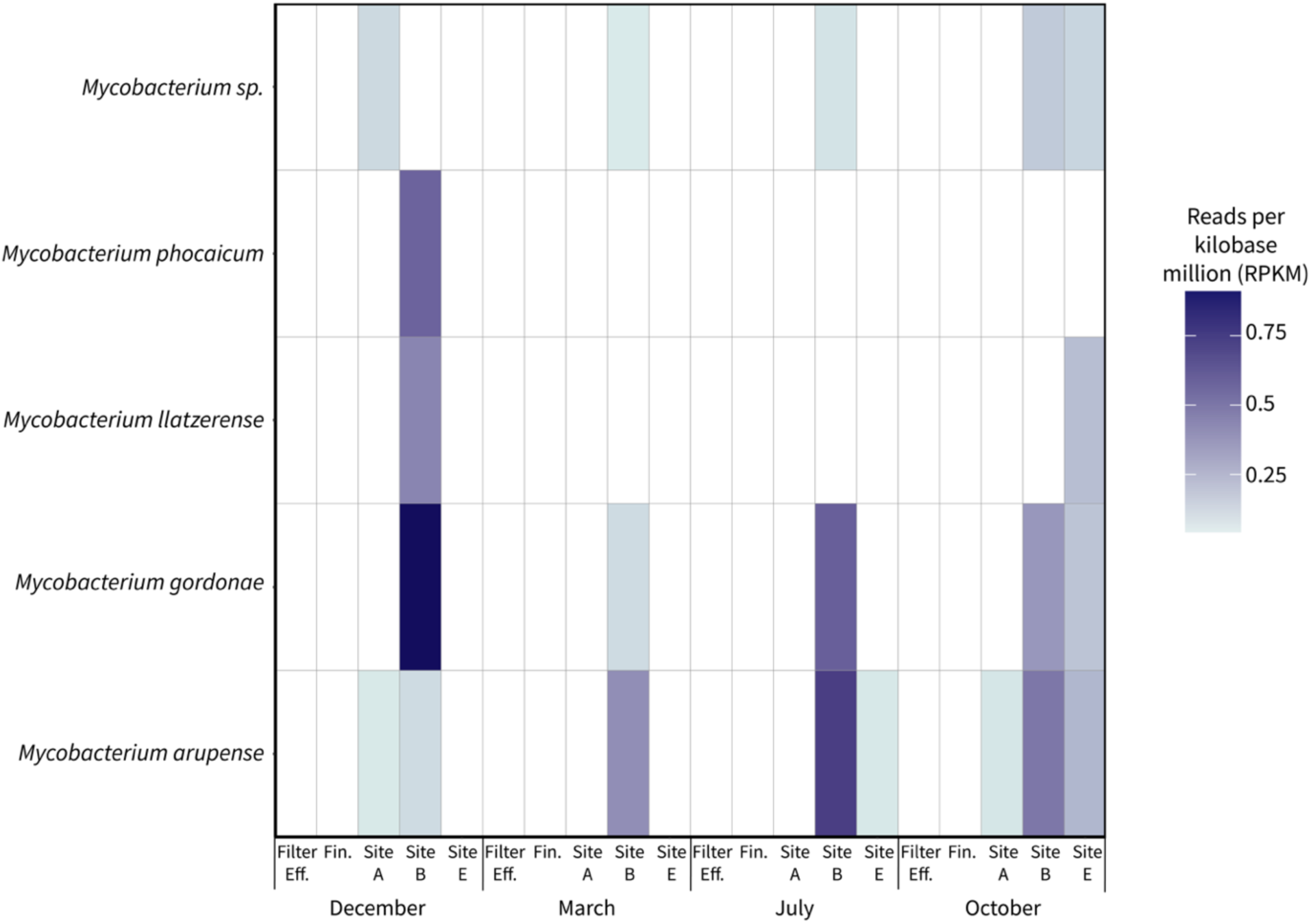
Metagenome-assembled genomes (MAGs) identified as *Mycobacterium*. Relative abundances of reads mapping to the NTM MAGs are represented by reads per kilobase million (RPKM). Filter Eff.: Filter Effluent; Fin.: Finished Water, Site A: Site A full flush sample, Site B: Site B full flush sample, Site E: Site E full flush sample.

The highest relative abundances for all the MAGs except for the *Mycobacterium* sp. MAG occurred in the Site B full flush samples. However, the months with the highest observed relative abundances varied, with the highest relative abundances for the *M. phocaicum*, *M. llatzerense*, and *M. gordonae* MAGs occurring in December, while the highest relative abundance of the *M. arupense* MAG occurred in July. The December Site B full flush sample was also one of only two samples in which four of the five MAGs were detected. The other sample in which four of the MAGs were detected was the October Site E full flush sample. While most of the MAGs were detected across seasons, *M. phocaicum* was only detected in December and *M. llatzerense* was only detected in December and October.

### Physicochemical and HPC analyses

Water sample temperatures fluctuated with the season, with temperatures ranging from less than 5°C in February 2020 to approximately 23°C in August 2020 (Figure S3). Although the Well-Source temperatures were relatively stable over the year (median: 11.9 ± 0.6°C, n=9), Well-Plant temperature trends were influenced by season (median: 18.0 ± 5.0°C, n=9). pH values did not vary substantially at any of the sampling locations over the sampling campaign (standard deviations ranged from <0.1 - 0.2; Figure S4). Median pH values were 7.3 in the well samples (Well-Source and Well-Plant; n=18), 8.1 in the river samples (River-Source and River-Plant; n=12), and 9.3 across all other sampling locations (n=175). Turbidities in the pre-filtration samples were typically greater than one nephelometric turbidity unit (NTU), with the highest turbidities occurring in the well water samples (median: 18.0 NTU, n=18) and the WTP ozone effluent (median:7.0 NTU, n=9; Figure S5). Median turbidity values for samples post-filtration were less than 0.3 NTU at all locations, except the first draw samples from Site E, where the median turbidity was 0.5 NTU (n=10). Monochloramine concentrations in the finished water were approximately 3.0 mg/L across all sampling events (median: 3.0 mg/L as Cl_2_, n=11; Figure 4). Building water sample monochloramine concentrations varied by location, sample type, and month. Of the first draw building water samples (median: 2.42 mg/L as Cl_2_, n=47), the lowest monochloramine concentrations were observed in the May and June samples at Site B, which were near or below the LOD of 0.04 mg/L as Cl_2_.

**Figure 4.**
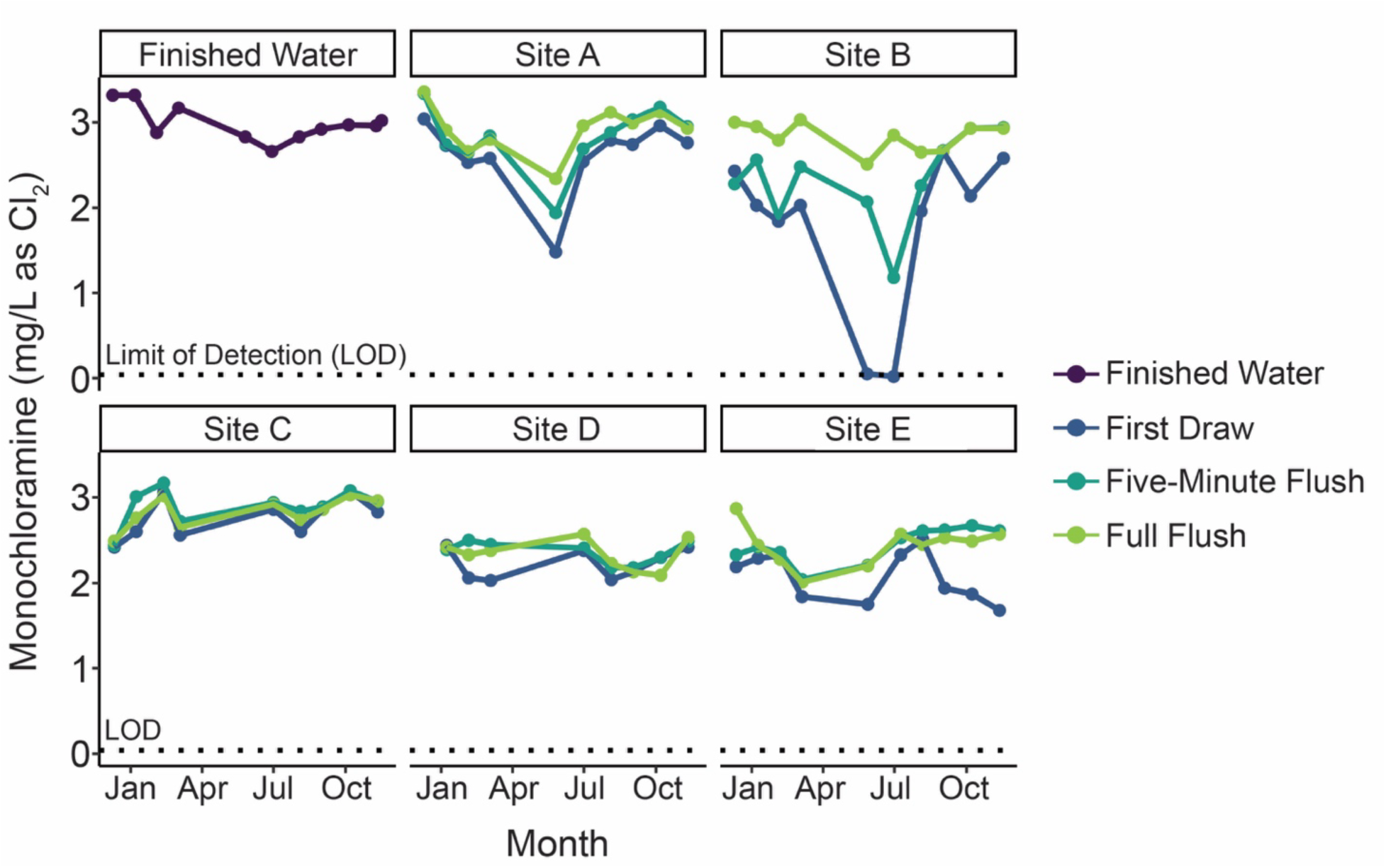
Monochloramine concentrations in the finished water and the five distribution system sites (Sites A – E). Samples at the distribution system sites included first draw, five-minute flush, and full flush. The dotted line indicates the limit of detection (LOD, 0.04 mg/L as Cl_2_). Source: Based on data from Dowdell et al., 2022 (43).

HPC results were generally low after secondary disinfection, except in some first draw samples (Figure S6). River water HPC results ranged from 1.5×10^3^ to 6.3×10^3^ CFU/mL (median: 3.9×10^3^ CFU/mL, n=7). HPC results were lower in the well water samples, which ranged from <1 CFU/mL to 8.3×10^2^ CFU/mL (median: 6.4×10^2^ CFU/mL, n=16). Notably, the HPC results were significantly higher in the Well–Plant samples compared to the Well–Source samples (p<0.05), indicating bacterial growth during transmission of the well water to the WTP. Ozone effluent HPC results ranged from 5 to 1×10^3^ CFU/mL (median: 1.2×10^2^ CFU/mL, n=9). Filter effluent HPC results were generally higher than in the ozone effluent, ranging from 18 to 2.6×10^4^ CFU/mL (median: 1.0×10^4^ CFU/mL, n=9). HPC results in the finished water ranged from 1 to 30 CFU/mL (median: 4 CFU/mL, n=11). In the distribution system, the full flush samples had the lowest HPC results overall (median: 8 CFU/mL, n=45) while the first draw samples were the highest (median: 1.7×10^2^ CFU/mL, n=45). However, median HPC results for first draw samples varied by site, with median results at Sites A (4.0×10^3^ CFU/mL, n=9), B (4.4×10^3^ CFU/mL, n=10), and E (median: 1.8×10^2^ CFU/mL, n=10) being at least one order of magnitude higher than the median HPC results at Sites C (41 CFU/mL, n=8) and D (4 CFU/mL, n=8).

### Impacts of decreased water use and flushing on physiochemical parameters and NTM

The COVID-19 pandemic occurred during sampling, resulting in lower than usual water usage in several distribution system monitoring locations beginning in mid-March 2020 (Figure 5). At Site A, the monthly water use in March, April, and May 2020 was at least 46% lower than in 2019, returning to values similar to 2019 beginning in June 2020. At Site B, March 2020 water use decreased to 48% of 2019 use, with the lowest water use in April and May 2020, both of which were at least 90% less than the monthly water use in 2019. At Site B, water use rebounded beginning in July 2020. At Site E, monthly water use in 2020 was below 2019 levels from March through May 2020, but then was substantially higher in the remaining months of 2020.

**Figure 5.**
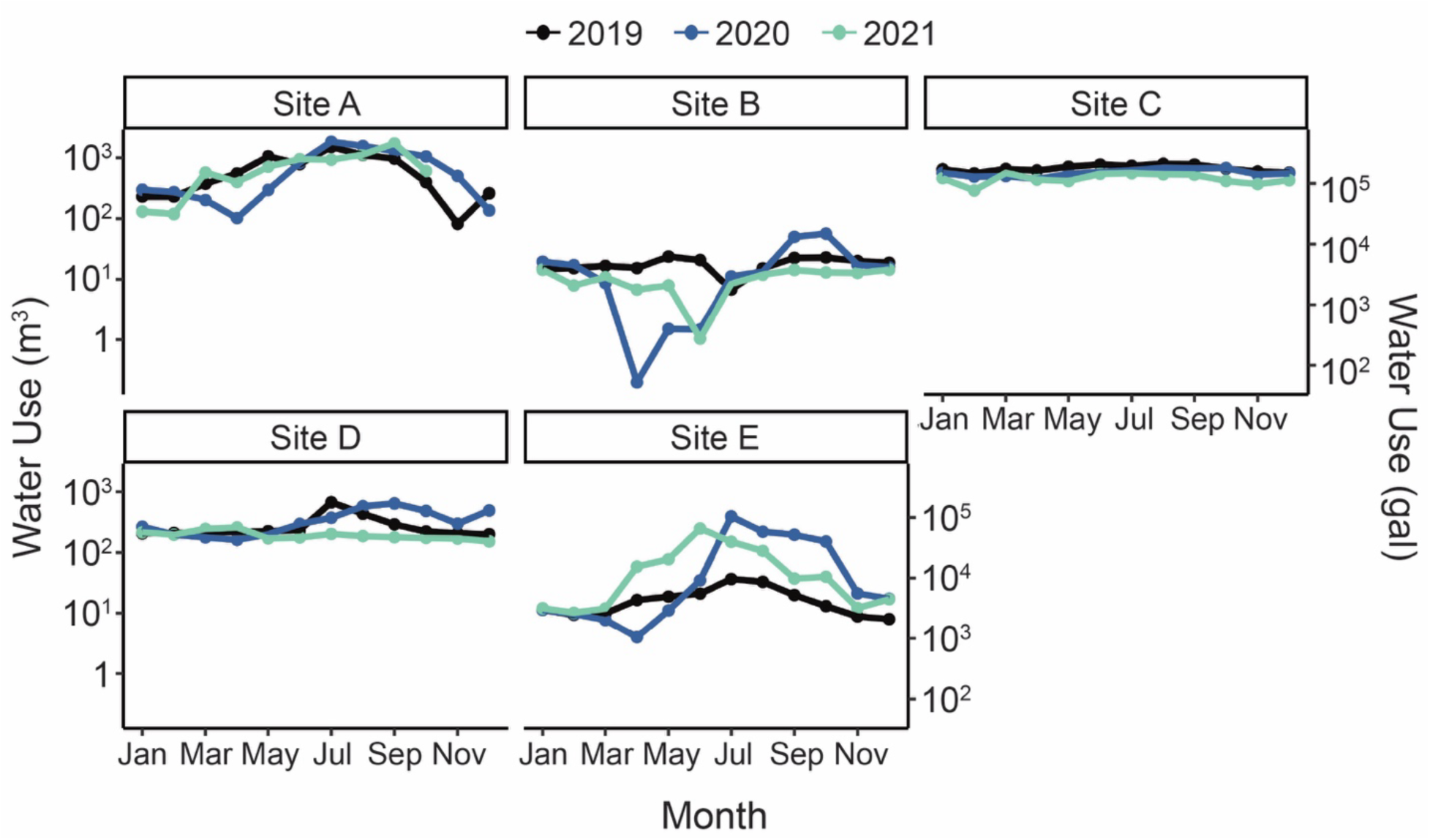
Monthly building water usage in the years 2019, 2020, and 2021. Units are cubic meters (left y-axis) and gallons (right y-axis).

The low water use at Sites A, B, and E, was associated with decreased disinfectant residual (Figure 4). Specifically, the monochloramine residuals in the first draw samples from Sites A and B in May 2020 were at least 40% lower than the monochloramine residuals in the March samples, which were collected prior to building closures. At Site B, first draw monochloramine residuals were near or below the detection limit (0.04 mg/L) in both May and July. At Sites A and B, first draw samples during the low water use period also had higher turbidities than the first draw samples once water use increased to normal levels (Figure S5).

Tap flushing improved water quality parameters at several building plumbing locations, though the degree of improvement and amount of flush time required for improved conditions varied by building and parameter. Flushing significantly reduced NTM qPCR results at Sites A and B (first draw versus full flush, p<0.05) but did not significantly reduce NTM qPCR results at Site C, D, and E. The flush duration required to reduce NTM qPCR results at Sites A and B varied. At Site A, the NTM concentrations in the five-minute flush and full flush were similar, suggesting that distribution system water was reached after five minutes flushing, or that building plumbing NTM levels were similar to distribution system levels. In contrast, flushing beyond five minutes reduced NTM gene copy concentrations by an average of 0.7 log at Site B.

### Influence of water age and physicochemical parameters on NTM

To investigate the influence of distribution on NTM and physicochemical parameters, correlation analysis was performed. The correlation analysis included temperature, pH, turbidity, monochloramine concentration, estimated water age at the site, and the qPCR results for the full flush samples. As this correlation analysis was focused on distribution system water age, first draw and five-minute samples were not included. The results showed that estimated water age had a strong negative correlation with monochloramine concentration (Kendall’s tau: −0.50, p<0.01; Figure S7). Estimated water age was also positively correlated with turbidity, though the association was weaker (Kendall’s tau: 0.24, p>0.05). The total NTM concentration was not strongly associated with any of the water quality parameters, though a weak positive correlation was observed with pH (Kendall’s tau: 0.23, p>0.05).

## Discussion

This study is unique in that monitoring had started several months prior to the COVID-19 pandemic, which allowed for evaluation of how decreased water use in buildings impacted general water quality parameters and NTM. At Sites A and B, where buildings were closed beginning in mid-March 2020, lower water qualities were observed in first draw samples, including turbidities as high as 4.6 NTU, HPC results as high as 8.0×10^4^ CFU/mL, and one sample without a measurable monochloramine residual. In addition to general water quality impacts, the concentrations of NTM gene copies were also higher, with values as high as 4.3×10^7^ gc/L. These NTM concentrations were similar to those reported in another study that investigated the impact of pandemic-related building closures on NTM using qPCR (16). As the buildings re-opened and water use increased in late summer and fall 2020, the NTM concentrations and physicochemical parameters, including monochloramine residual, generally improved. In Sites C and D, which remained fully open during the pandemic lockdown, water use was maintained as usual. This continuation of similar water use patterns was reflected in the consistency of the NTM concentration and monochloramine residuals across the year of sampling. Although most studies investigating the impacts of COVID-19 pandemic-related building closures focused on other parameters, such as metals or *L. pneumophila* (32–34, 80), this study shows the importance of also considering the impact of low water use on NTM occurrence.

By sampling from drinking water sources to taps in buildings in the distribution system, this study investigated how NTM populations change through water treatment and delivery. In well water, NTM gene copy concentrations changed even prior to treatment, increasing during conveyance from the well to the WTP. Transmission of well water to the WTP also significantly increased water temperatures (p<0.05; Figure S3) and HPC results (p<0.05, Figure S6) in the Well-Plant samples compared to the Well-Source samples. As a result, the temperature curve for the Well-Plant samples is more similar to that of the surface water samples than the Well-Source samples (Figure S3). While ozonation effectively reduced NTM gene copy concentrations to below the LOQ, these concentrations rebounded after filtration (median:10^4^ gc/L) and only decreased after secondary disinfection in certain months. These findings support previous studies, which observed that NTM concentrations increased after filtration (25, 81). Additionally, these results are consistent with the importance of filtration in shaping bacterial communities in the finished water (82).

Plate culture analyses indicated that the WTP samples were dominated by non-NTM, including spore-forming bacteria and microorganisms that could not be identified. The first draw samples from Sites A and B and the full flush sample from Site B also yielded either all or a majority of non-NTM isolates. The WTP filter effluent was the first location where NTM isolates were recovered from culture plates and, except for Site A and B first draw samples, the building plumbing sample isolates were at least 75% NTM. The low recovery of NTM isolates prior to filtration is likely due to high concentrations of non-NTM microorganisms in those samples. In previous studies focused on NTM quantification in source waters or varied water types, antibiotics were generally added to the NTM culture media in addition to using decontamination strategies (23, 38). Therefore, only using decontamination, as was done in this study, was insufficient for the recovery of NTM in certain samples with high concentrations of non-target microorganisms. However, the selected culture method and similar methods utilizing cetylpyridinium chloride (CPC), a common biocide, as a pre-treatment have been commonly used to quantify NTM in drinking water samples, including building water samples (22, 30, 35), and the method selected in this study performed well at all sites post-disinfection except for the first draw samples from Sites A and B.

The occurrence of specific species of NTM varied by site, with *M. aurum* only isolated from the WTP filter effluent, *M. llatzerense* only recovered from Site A, *M. avium* only recovered from Sites D and E, and *M. arupense, M. asiaticum*, and *M. gordonae* only isolated from Site E. The finding that *M. avium* was only recovered from the sites with the highest water ages supports the results of a previous study, which reported that the relative abundance of *M. avium* increased with water age (14). Other NTM species, including *M. chelonae* complex, *M. mucogenicum*/*phocaicum* group, and *M. franklinii* were isolated from across sites with varying water ages. Overall, the NTM isolates were predominantly *M. chelonae* complex (43%, n=84) and *M. mucogenicum/phocaicum* group (34%, n=65). A recent study investigating NTM species in chloraminated distribution systems also reported that *M. mucogenicum/phocaicum* group was the most common species recovered, though in that study the proportion was higher at 76% of the isolates (21).

Among the NTM isolates recovered in this study, *M. avium* and *M. chelonae* complex are most associated with human infection (83). While *M. avium* was only isolated from the full flush samples from Site D (n=5) and Site E (n=3), *M. chelonae* complex was the most commonly recovered isolate, representing 26% (n=84) of all isolates and 43% of NTM isolates. Though not as commonly associated with disease as *M. avium* and *M. chelonae*, *M. mucogenicum* and *M. peregrinum* have also been reported to cause human infections (57, 84, 85). The prevalence of clinically-relevant NTM species other than *M. avium* emphasizes the importance of monitoring for additional NTM species, such as *M. abscessus* and *M. chelonae*, when investigating NTM occurrence in drinking water. Studies have also reported geographical differences in the NTM that account for the majority of infections (49, 56), which should be considered when selecting species-specific methods for NTM quantification.

Metagenomic analysis detected four species of NTM in the samples analyzed: *M. phocaicum*, *M. gordonae*, *M. llatzerense*, and *M. arupense*, all of which were also recovered by plate culture. However, there were differences in where these NTM species were detected when using plate culture versus metagenomic analysis. *M. phocaicum*, which was detected in the December Site B full flush sample by metagenomic analysis, was isolated by plate culture from the filter effluent and all distribution system sampling sites. *M. arupense*, which was detected at Sites A, B, and E by metagenomic analysis, was only recovered from Site E with plate culture. Similarly, *M. gordonae* was detected in Site B and E samples by metagenomic analysis but only in Site E by plate culture, and *M. llatzerense* was detected in Sites B and E using metagenomic analysis but only in Site A by plate culture. Therefore, both methods detected certain species that were not identified using the other method at particular sites. Possible reasons for this disagreement include that the sequencing depth may have been insufficient to detect rarer species, that not all microbial community diversity was captured during binning, that the smaller sample volumes associated with the plate culture method may have prevented detection of rarer species, and that the plate culture method may have de-selected for certain species. These results highlight the benefits of using both cultivation-dependent and -independent methods, as it allows for a more complete understanding of NTM occurrence.

Although the culture-based method resulted in the identification of a larger number of NTM species, there are known biases associated with NTM plate culture methods. The pre-treatment step, for example, has been shown to impact species and strains of NTM differently, with one study reporting that a dose of 0.1% CPC for 30 minutes reduced culturable water-grown *M. avium A5* by approximately 80% but only reduced *M. avium Va14 (T)* by 20% (36). Differences in susceptibility of NTM species to pre-treatments by growth category (rapid-growing or slow-growing) have also been observed, with one study reporting that rapidly growing NTM species such as *M. fortuitum*, *M. abscessus*, and *M. peregrinum* may be more sensitive to decontamination (37). However, it is difficult to balance the need to reduce colony formation by non-NTM with recovery of target species. A new NTM plate culture medium that can be used without sample pre-treatment has been described recently (86). This method may overcome some of the limitations of existing methods, though additional comparisons are needed to validate its selectivity (86, 87).

Sampling over the course of a year allowed for investigation of the impacts of temperature and season on NTM populations. While strong seasonal trends were not observed, the NTM gene copy concentrations in full flush samples across all sites were lowest in November (median: 2.9×10^3^ gc/L, n=5) and highest in March (3.3×10^4^ gc/L, n=5). The impact of season could not be addressed for the first flush and five-minute flush samples due to the changes in water use patterns at several of the sites due to the COVID-19 pandemic. The impact of season on NTM species occurrence also could not be determined based on the MADLI-TOF MS results due to the low numbers of isolates analyzed per site per month and the strong influence of site on NTM species. The metagenomic analysis found the highest relative abundances of NTM MAGs in December and July. Overall, low water use appeared to be the driving factor of NTM concentration in building plumbing rather than season. While few studies have investigated the impact of season on NTM concentrations, one previous study reported higher concentrations of NTM gene copies in a distribution system during the summer compared to the winter (88). However, another study reported significantly higher concentrations of NTM in distribution systems in winter and spring, which it attributed to lower water use (19). Due to COVID-19 pandemic-associated building closures during this study, it is difficult to determine whether the generally higher NTM concentrations observed in the summer are linked to season or were the result of shifts in water use patterns. Additional studies are needed to further investigate the potential impacts of both water use and season on NTM in drinking water distribution systems.

## Methods

### Source water and treatment plant

The full-scale WTP sampled in this study has been described previously (15, 27, 82). Briefly, the City of Ann Arbor WTP is a 50 million gallon per day (1.9×10^5^ m^3^/day) facility in Ann Arbor Michigan, USA that treats groundwater and river water, which are blended at a ratio of approximately 1:6, using two-stage excess lime softening, coagulation, flocculation, sedimentation, ozonation, biological filtration, and monochloramine disinfection (3 mg/L as Cl_2_). Ozonation is used to achieve 0.5-log removal of *Cryptosporidium*. Typical ozone concentration × time (CT) values range from 0.3 to 1 mg/L. The biological filters contain either dual media sand-granular activated carbon (GAC) or GAC alone. In June 2020, the WTP began using an ultraviolet (UV) disinfection system intermittently, which was intended to provide additional *Cryptosporidium* inactivation when the plant is operated with single-stage lime softening during maintenance periods. The UV system was in operation during the June, August, and September sampling events.

### Sample collection

Samples were collected approximately monthly from December 2019 through November 2020. Samples were not collected due to the COVID-19 pandemic in April 2020, and only a partial sampling event (finished water reservoir [Finished Water] and three buildings in the distribution system [Sites A, B, and E]) occurred in May 2020 due to building access restrictions. Samples were collected using sterile polypropylene bottles. All source water and treatment plant samples except the Ozone Effluent samples were either collected from the sampling taps that were continuously flowing (River-Plant, Well-Plant, and Finished Water) or were collected after flushing the taps for at least five minutes (River-Source, Well-Source, and Filter Effluent). Source water samples were collected from sampling taps at the source water pump stations (River–Source and Well–Source) as well as from the sampling taps at the WTP used for compliance monitoring (River–Plant and Well–Plant). River sample volumes ranged from 2 to 4 L and well samples ranged from 10 to 15 L. The Ozone Effluent samples were collected from a clearwell after the ozone contactors and prior to filtration. Ozone Effluent sample volumes ranged from 10 to 20 L. Filter Effluent samples were the combined effluents of six filters and ranged in volume from 10 to 20 L. Finished Water samples were collected from a sampling tap in the laboratory of the WTP used for compliance monitoring.

Cold water samples were collected from sinks in five buildings, Sites A – E, receiving water from the WTP via the distribution system (Table S1). These locations are a subset of the 13 used for distribution system compliance monitoring. Sites were selected to capture the full range of estimated water ages at compliance sites, as determined using an EPANET distribution system model (89). Sites were ordered based on estimated water age and selected to capture a range of water quality characteristics based on historical data provided by the WTP, which included concentrations of total chlorine and nitrogen species, and HPC. Additional details on the distribution sampling locations are included in Table S1. Three samples were collected from each distribution sampling location: 1) a 2 L first draw sample, representing the water in the fixture and building plumbing line supplying the fixture, 2) a 10 L five-minute flush sample, meant to represent the typical monitoring sample collected by the utility, and 3) a 10 to 20 L fully flushed sample, meant to represent distribution system water at the location. To mimic the procedure for compliance sampling, the tap was not disinfected for the first draw samples but was sprayed with a 10% bleach solution prior to collection of the five-minute flush and full flush samples. For the full flush samples, temperature and pH were monitored and samples were collected when successive readings stabilized.

### Physicochemical analyses

Temperature, pH, conductivity, and total dissolved solids (TDS) were analyzed immediately on-site using a combination probe (Hanna Instruments HI98121, Smithfield, RI, USA). Total chlorine, free chlorine, and monochloramine were also analyzed on-site using a portable colorimeter (Hach DR900, Loveland, CO, USA) and powder pillows (Hach Methods 10250, 10245, and 10200). Samples for total organic carbon (TOC) were transferred to carbon-free glass vials and acidified with 85% phosphoric acid. TOC samples were analyzed with the non-purgeable organic carbon (NPOC) method using a Shimadzu TOC analyzer (TOC-V, Japan). Turbidity was measured using a benchtop turbidimeter (Hach TU5200).

### DNA collection and extraction

Water samples were filtered onto sterile 0.22 μm copolymer cartridge filters (EDM Millipore, USA, cat. no. SVGPL10RC) using sterile tubing (Masterflex, USA) and peristaltic pumps. Filters were then placed in sterile bags. Samples collected from December 2019 through March 2020 were filtered in the field and flash frozen using dry ice and ethanol, then transported to the laboratory and frozen at −80°C. Due to COVID-19 pandemic restrictions, samples collected after March 2020 were transported in coolers with cold packs to the laboratory for filtration and were placed in the −80°C freezer rather than flash freezing with dry ice.

DNA was extracted using a modified form of the QIAGEN DNeasy PowerWater kit (QIAGEN, Hilden, DEU, cat. no. 14900-100-NF) method, which includes additional enzymatic (Proteinase K and lysozyme) and chemical (chloroform-isoamyl 24:1) lysis steps (90). DNA purity was measured using a Nanodrop 1.0 spectrophotometer (Thermo Fisher Scientific, Waltham, MA, USA) and DNA concentrations were measured using the Qubit™ dsDNA High-Sensitivity assay kit (Thermo Fisher Scientific, cat. no. Q32851).

### NTM and HPC culture methods

Samples for HPC analyses were transferred to 100 mL sterile vessels containing excess sodium thiosulfate and analyzed using the pour-plate method (SM 9215-B-2000) (91). NTM plate culture was performed for all sampling events except February, May, and August 2020. Sample volumes for NTM plate culture ranged from 3 mL to 1 L and were adjusted each month based on colony counts for the previous month in an effort to maximize the number of plates that yielded 1 to 300 colonies (Table S2). Samples (n=148) were processed in duplicate except when insufficient volume was available to process in duplicate (occurred in 13 cases, all of which were first draw samples with limited volume). Filter concentration followed by pre-treatment with CPC was employed. Briefly, samples were transferred to sterile glass bottles and dosed with a sterile 2% CPC solution to a final concentration of 0.04%. Samples were incubated at room temperature for 30 minutes, then immediately filtered onto 0.45 μm filters (Thermo Fisher Scientific, cat. no. 09-719-555) using sterile glass filtration bowls and pedestals and a vacuum filtration manifold (EDM Millipore, USA) in a fume hood. After filtration, the filters were rinsed with 50 mL of sterile ultrapure water (Invitrogen™ UltraPure™ DNase/RNase-Free Distilled Water, Thermo Fisher Scientific, cat. no. 10977015) to remove residual CPC. Filters were plated onto sterile Middlebrook 7H11 agar (Thermo Fisher Scientific, cat. no. R454002) supplemented with oleic albumin dextrose catalase (OADC, BBL™ Middlebrook OADC Enrichment, BD Biosciences, Franklin Lakes, NJ, USA, cat. no. 212351) using sterile tweezers in a biological safety cabinet. Plates were allowed to dry in the biological safety cabinet, then sealed with Parafilm (Bemis Company, USA), inverted, and placed in loosely sealed plastic bags. Plates were incubated in the dark at 35°C for at least one month and observed at least once per week. Controls were included for at least every sampling day and included plate controls (agar plates only, n=24) and filtration controls (50 mL of ultrapure water filtered and plated, n=26).

Plates were classified into four groups, either negative (no colonies), positive and quantifiable (CFU between 1 and 300), too numerous to count (TNTC; CFU greater than 300), or no count (NC). The NC category represented plates that contained spreader colonies or other colonies that grew to overtake the plates, preventing accurate counting of colonies, which may have originated in the samples or may have been the result of contamination. Colony counts from the duplicate plates were averaged to obtain a plate count for each sample. For calculations and plotting, TNTC plate counts were set to 300 CFU and negative plates were set to 0.5 CFU. Excluding controls, a total of 284 plates were processed for 148 samples. Of these, 33 plates (12%) were TNTC, 44 (15%) were negative, and 20 (7%) fell in the NC category, leaving 187 plates (66%) from 107 (72%) samples that were counted. Only four of the 50 controls showed contamination. For all plates that were counted, at least 10% of the colonies were picked for confirmation testing. Colonies were selected to cover a range of morphologies and growth characteristics. Colonies were picked in a biological safety cabinet using sterile pipette tips. Biomass was then transferred to sterile 0.6 mL tubes containing 50 μL of ultrapure water and frozen at −80°C.

### NTM isolate protein extraction MALDI-TOF MS analysis

Colonies (n=391) from the NTM culture plates from the December, January, March, June, and October sampling events were analyzed using MALDI-TOF MS. Due to the number of isolates, only isolates from first draw and full flush samples were selected for analysis. Briefly, colonies were thawed and used to inoculate 5 mL tubes containing Middlebrook 7H9 with Tween® 80 (Hardy Diagnostics, Santa Maria, CA, USA, cat. no. C62) and incubated at 35°C. Once growth was observed, isolates were streaked onto Middlebrook 7H11 with OADC and incubated at 35°C. Isolates that failed to grow on Middlebrook 7H11 with OADC at 35°C were streaked onto a Middlebrook 7H11 with OADC and incubated at 32°C and streaked onto an LB agar and incubated at 35°C. Seventy of the isolates (18%), from 25 samples, failed to grow despite the use of multiple temperatures and media and therefore could not be analyzed using MALDI-TOF MS.

For the protein extraction, each isolate was extracted in duplicate by picking one μL loopfuls of colonies from the culture plates and transferring them to two 2 mL tubes containing 200 μL of 0.5 mm zirconium/silica beads and 500 μL of 70% ethanol. Beads were heat-treated prior to use by incubating in a 450°C oven for at least three hours. Tubes were vortexed for 15 minutes, then centrifuged at 10,000xg for two minutes. The supernatant was discarded, and tubes were allowed to dry in a biosafety cabinet. Once dry, tubes were transferred to a fume hood, where 20 μL of 70% formic acid was added to each tube. Tubes were vortexed and incubated for five minutes, then 20 μL of acetonitrile was added. Tubes were centrifuged again at 10,000xg for two minutes and the supernatant was used for MALDI-TOF MS analysis within four hours of extraction. Each protein extraction session included a negative control (an empty bead tube), a non-NTM positive control (colonies of *Escherichia coli*), and an NTM positive control (colonies of *Mycobacterium smegmatis*).

MALDI-TOF MS analysis was performed at the University of Michigan Microbiome Core using a Bruker MALDI Biotyper® sirius system (Bruker Daltonics, Billerica, MA, USA) and a 96 target brushed steel plate. Each MALDI-TOF MS plate analysis began with analyzing two targets of Bacterial Test Standard (BTS, Bruker Daltonics, Billerica, MA, USA, cat. No. 8255343) to ensure the system was functioning properly. All extracts were spotted in duplicate and spectra were analyzed using both the standard instrument library and the *Mycobacterium* genus-specific Mycobacteria RUO Library (Bruker Daltonics). BTS scores were required to be above 2.0 and *M. smegmatis* positive control results were required to be above 1.7. The maximum score between the duplicate extractions and duplicate spots (four results per isolate) was used for identification. Minimum scores for identification followed manufacturer guidance; NTM isolates were required to score at least 1.6 to be identified to the species or complex level, while non-NTM isolates were required to score at least 1.7 (68). Per the manufacturer, NTM isolates with scores less than 1.6 were not identifiable, isolates with scores between 1.60 and 1.79 were “low confidence” identifications, and isolates with scores of at least 1.8 were considered “high confidence” identifications. Score results by genus and species are provided in Table S3. Due to the inability of the method to distinguish between *M. chelonae*, *Mycobacterium stephanolepidis*, and *Mycobacterium salmoniphilum*, these species are reported as *M. chelonae* complex. The method also cannot distinguish between *M. mucogenicum* and *M. phocaicum*, so matches are reported as *M. mucogenicum/phocaicum* group. The MALDI-TOF MS results for isolates were used to adjust the presumptive NTM CFU/mL by multiplying the presumptive CFU/mL by the percentage of the isolates from the sample that were identified as NTM.

### NTM qPCR

qPCR analysis followed the guidance of the Minimum Information for Publication of Quantitative Real-Time PCR Experiments (MIQE) and Environmental Microbiology Minimum Information Guidelines (EMMI) (92, 93). Gene copies of the *Mycobacterium atpE* gene were quantified using the Radomski et al. assay (FatpE: 5’-CGGYGCCGGTATCGGYGA-3’; RatpE: 5’-CGAAGACGAACARSGCCAT-3’) (41), generating an approximately 164 base pair amplicon, with the modification that EvaGreen was used instead of probe chemistry (15). qPCR was performed using 96-well plates containing 10 μL reactions composed of 5 μL of master mix (Biotium FastEvaGreen 2x, final concentration 1x, cat. no. 31003), 0.5 μL each 10 μM forward and reverse primers (final concentration of 0.5 μM, Integrated DNA Technologies [IDT], Coralville, IA, USA), 0.25 mL of 25 mg/mL bovine serum albumin (Thermo Fisher Scientific, cat. no. AM2616, final concentration 0.625 mg/mL), 2.75 μL of ultrapure water and 1 μL of template. Samples were analyzed using a real-time PCR system (Applied Biosciences QuantStudio 3, Thermo Fisher Scientific) using the following cycling conditions: 95°C for 5 minutes, 35 cycles of 95°C for 20 seconds, 59.6°C for 30 seconds, and 72°C for 30 seconds. Melt curve analysis was performed for all plates. Standard curves were prepared using synthetic DNA (gBlock, IDT, USA), consisting of the amplicon with 30 base pairs of neutral adaptors on both ends. Standard curves were included on each 96-well plate. Negative controls with ultrapure water in place of template were also included on each 96-well plate. The LOD (41 gc/rx) and LOQ (41 gc/rxn) were determined using serial dilutions of the standards. Samples were diluted to reduce inhibition, with dilution factors ranging from undiluted to 1:110. Inhibition was assessed on the majority of samples (88%, n=141) by analyzing samples at two dilution levels and comparing the actual Cq to the theoretical Cq. Thresholds were automatically set by the qPCR instrument. All plates were required to meet a minimum efficiency of 85% and R^2^ standard curve results of at least 0.98. Results were converted from gc/rxn to gc/L using the volume of template per reaction (1 μL), the dilution factor, the DNA extraction elution volume (100 μL), and the filtered sample volume.

### Metagenomic analysis

The filter effluent, finished water, and full flush samples from distribution system Sites A, B, and E from the December, March, June, and October sampling events (n=20) and three negative controls (pooled filter controls, filtration controls, and DNA extraction controls) were submitted for shotgun metagenomic sequencing. Library preparation, sequencing, and de-multiplexing were performed by the University of Michigan Advanced Genomics Core. Paired-end sequence libraries were prepared using the NEBNext® Ultra™ II FS DNA Library Prep Kit for Illumina (New England BioLabs Inc., Ipswich, MA, USA, cat. no. E7805S). Sequencing was performed using the NovaSeq 6000 system and an SP flow cell with 500 cycles, producing 250 nucleotide paired-end reads.

Bioinformatic analysis of reads followed the pipeline described in Vosloo et al. (94). Briefly, pre-processing was performed using fastp (v0.20.0) (95) to remove adaptors and perform initial quality filtering. Reads mapping to the UniVec_Core database (ftp://ftp.ncbi.nlm.nih.gov/pub/UniVec/) and negative controls were removed as potential contamination. The cleaned reads were pooled and *de novo* co-assembly into contigs was performed using metaSPAdes (v.3.13.1) with kmer sizes of 21, 33, 55, 77, 99, and 127 (96). Contigs less than 1 kilobase pair were removed using seqtk (https://github.com/lh3/seqtk) and redundant contigs were removed using the *dedupe* function of BBTools (v38.76) (97). The quality of the assembly was determined using QUAST (v.5.0.2) (98) and assembly validation was performed by calculating the mapping rate of processed reads to contigs. Binning was performed using Anvi’o (v6.1) workflow for the analysis and visualization of omics data. Binning algorithms CONCOCT (v.1.1.1), MetaBAT (v.2.12.1), and MaxBin (v.2.2.4) (99–102) were used together with the bin aggregation software DAS Tools (v.1.1.0) to select the highest quality bins with the least redundancy (102). Bin statistics, including bin size, GC content, and number of contigs, were determined using the summarize function in Anvi’o. Further bin quality estimates, including completeness, redundancy, and strain heterogeneity, were determined using CheckM (v 1.0.18) (104). To improve bin quality, bins with at least 50% completeness were reassembled using metaSPAdes with the same kmer sizes previously used. Re-binning was performed using the same binning strategy and manually curated using Anvi’o. Duplicate bins were dereplicated using dRep (v2.6.2) and clustered into species-level representative genomes using a 95% average nucleotide identity (105). The species-level representative genomes were classified using the Genome Taxonomy Dataset Toolkit (GTDTk, v0.3.2) (106) and final bin statistics and quality were determined as described previously. CoverM (v0.4.0) was used to calculate coverage of the MAGs across samples with RPKM as a metric for relative abundance (107). The covered_bases parameter of coverM was also used to calculate the number of bases covered by one or more reads at a coverage threshold of 25%, indicating that only bases with sequencing read coverage equal to or greater than 25% of the expected coverage depth were considered as covered.

### Data analysis

Data cleaning and analyses were performed using R (version 4.1.1) and R Studio (1.4.1717) (108, 109). R packages used for analyses include *ggplot, dplyr, lubridate, viridis, readxl, and stats*. For the qPCR data analysis and plotting, values less than the LOQ were set at one-half the LOQ. For physicochemical parameters, values less than the LOD were set at one-half the LOD. Statistical analyses were performed using the *stats* package. Differences between sample locations, sample types, and by season were calculated using the Wilcoxon signed-rank test using a significance threshold of 0.05. Correlation analysis and correlation plot were performed using the functions “cor” and “cor.test” from the R package *stats* and the R package *corrplot* using Kendall’s tau b.

## Acknowledgments

The authors gratefully acknowledge the staff of the City of Ann Arbor WTP and building owners and staff for their support of this work. The authors thank Ruqaiya Siddiqui, Michael Bachman, Stephanie Agozino, and Jennifer Furlong for supporting MALDI-TOF MS analysis and Bridget Hegarty for helpful discussions. The authors thank Ameet Pinto and Solize Vosloo for supporting the analysis of the metagenomics data. Funding was provided by the Blue Sky Initiative (College of Engineering, University of Michigan) and Water Research Foundation Project Number 4721. K.S.D was supported by a National Science Foundation Graduate Research Fellowship (grant number DGE-1256260) and a University of Michigan Rackham Predoctoral Fellowship. This research was supported by work performed by The University of Michigan Microbiome Core.

## Data Availability Statement

The raw sequencing data are available through the National Center for Biotechnology Information (NCBI) under BioProject No. PRJNA1081894.

## Supplemental Information

Supplementary materials are available in the Supplementary Information file.

